# The human pangenome reference reduces ancestry-related biases in somatic mutation detection

**DOI:** 10.64898/2026.03.30.715289

**Authors:** Chau V.K. Pham, Farida S.A. Abdelmalek, Tracy Hua, Erika Apel, Ava Bizjak, Ethan J. Schmidt, Kathleen E. Houlahan

## Abstract

Commonly used human reference genomes collapse extensive genetic variability into a single linear genome of which 70% is derived from one donor. These linear genomes fail to capture the full spectrum of genetic variation, which can lead to misalignment of sequencing reads particularly for individuals underrepresented by the linear reference genomes. To address this shortcoming, the Human Pangenome Reference Consortium released the first draft of the human pangenome reference, a graph-based reference that integrates diverse haplotypes. While the human pangenome reference has shown increased accuracy in detecting inherited DNA variants, it remains to be seen if the observed improvements extend to somatic mutation detection. Here, we systematically benchmarked somatic single nucleotide variant (SNV) detection leveraging the human pangenome in 30 whole exome sequenced bladder tumours with matched blood tissue of diverse ancestries. We found somatic SNV detection leveraging the human pangenome reference outperformed the linear reference, most notably in individuals of East Asian ancestry where we observed on average a 20% improvement in detection accuracy. Improvements to detection accuracy in individuals of European ancestry were marginal. The increase in accuracy was attributed to reduced germline contamination and reduced reference bias. Further, we demonstrate the pangenome increases SNV detection precision, mitigating the need for time and computationally expensive ensemble approaches that take the consensus across multiple tools. Finally, we demonstrate that the increased precision when aligned to the pangenome generalized to an additional 29 lung adenocarcinoma tumours, particularly for individuals of East Asian ancestry. These findings support adoption of the pangenome to improve somatic variant detection and reduce ancestry-related disparities.

## INTRODUCTION

Current genomics workflows designed to identify somatic mutations run the risk of potentiating bias towards individuals of European descent. This is in part because genomics workflows require alignment of sequencing reads to a reference genome. Commonly used human reference genomes collapse extensive genetic variability into a single linear genome of which 70% is derived from a single donor^1^. Unsurprisingly, these linear genomes fail to capture the full spectrum of genetic variation, which can lead to misalignment of sequencing reads particularly for individuals underrepresented by the linear reference genomes.

To address this shortcoming, the Human Pangenome Reference Consortium released the first draft of the human pangenome reference containing 47 phased, diploid assemblies from a diverse catalogue of individuals. Indeed, in contrast to previous studies, only 2% of the cohort were individuals of European ancestry, with a larger representation of individuals of African (51%), Caribbean/South American (34%) or Asian (13%) ancestry. The human pangenome reference improved detection of inherited single nucleotide variants by 34% and larger structural alterations by 104%^2^. These proof-of-principal studies have demonstrated the value of pangenome reference genomes in detecting inherited variation, but it remains to be seen if this benefit will translate to somatic mutation detection.

Accurate somatic mutation detection underpins effective biomarker discovery and clinical implementation. Miscalling somatic mutations can exacerbate health disparities, particularly when the likelihood of miscalling is linked to genetic ancestry. As a representative example, Nassar *et al*. found that quantification of the tumour mutation burden (TMB), an FDA-approved biomarker for the administration of the immune checkpoint inhibitor pembrolizumab, from tumour samples alone lead to inaccurate overclassification of tumours as TMB-high due to germline variants being miscalled as somatic^3^. This overclassification was particularly pronounced in individuals of African and Asian ancestry. These data emphasize how ancestry-related biases in genomics workflows can lead to biased clinical decision making.

To address if the incorporation of the human pangenome reference in somatic mutation detection workflows can mitigate ancestry-related biases, we systematically benchmarked somatic SNV detection leveraging the human pangenome in 30 whole exome sequenced bladder tumours with matched blood tissue from The Cancer Genome Atlas (TCGA)^4^. We selected bladder cancer because it is a highly point mutated tumour^5^ and has well documented differences in incidence, prognosis and somatic mutation landscape linked to genetic ancestry^6–8^. We focused on the exome reasoning that it represents the most well characterized region of the genome and, thus, would provide a lower-bound on the anticipated improvements expected. It also more closely reflects improvements expected during biomarker measurements in the clinic. We found somatic mutation detection leveraging the human pangenome reference outperformed the linear reference, most notably in individuals of East Asian ancestry where we observed on average a 20% improvement in detection accuracy. Improvements to detection accuracy in individuals of European ancestry were more marginal. The increase in accuracy was attributed to reduced germline contamination and reduced reference bias. The same increase in precision, primarily in individuals of East Asian ancestry, was replicated in a second cohort of 29 whole exome sequenced lung adenocarcinoma tumours. Taken together, these data motivate the use of the human pangenome reference into genomics workflows to ensure equitable and accurate somatic mutation detection.

## RESULTS

### The human pangenome improves sequence alignment

To benchmark somatic SNV detection leveraging the human pangenome, we selected 30 whole exome sequenced bladder tumours with matched blood tissue from TCGA^4^ (**Figure 1a; Supplementary Figure 1a**). To ensure our cohort was balanced across ancestries, we selected ten donors each of European, African and East Asian ancestry based on genetically inferred ancestry (**Supplementary Figure 1b-d**). DNA sequencing for both tumour and blood tissue was aligned to the linear GRCh38 reference genome (referred to as the linear reference) as well as CHM13-T2T pangenome^2^ (referred to as the pangenome reference), which includes both GRCh38 and T2T reference genomes (**Figure 1a; Supplementary Figure 1a**). While there was no significant difference in the total number of reads that mapped between the linear and pangenome references (P = 0.594; Wilcoxon Signed-Rank Test; **Supplementary Figure 2a-b)**, mapping to the human pangenome reference resulted in significantly more properly paired reads than the linear reference (fold-change (FC) = 1.02; P < 1.67x10^-11^; Wilcoxon Signed-Rank Test; **Supplementary Figure 2c-d**). Taken together, these data indicate mapping to the human pangenome reference results in improved sequencing alignment, even within the most well characterized regions of the genome.

**Figure 1.**
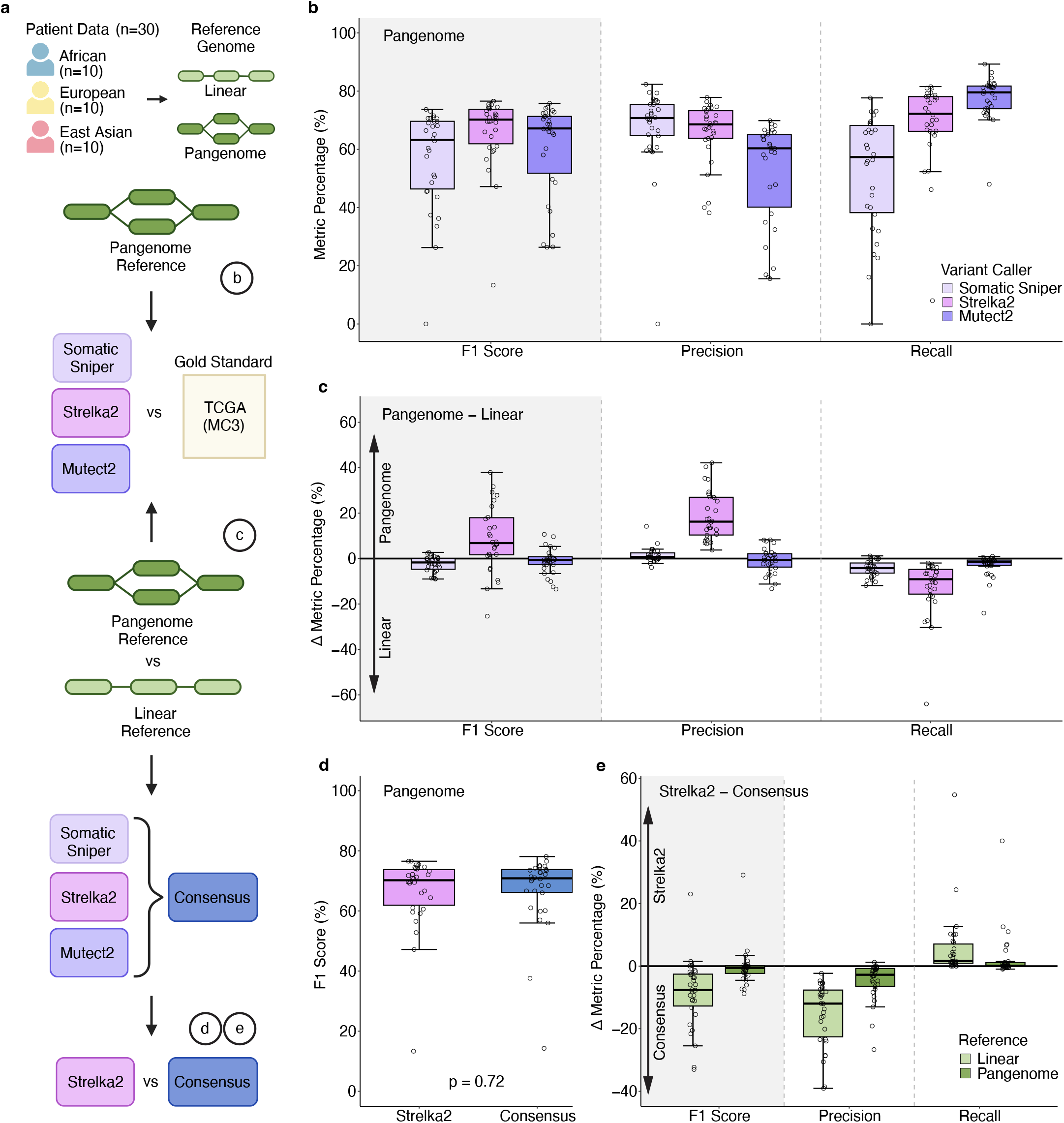
The human pangenome improves the precision of somatic mutation detection. **a)** Schematic overview of benchmarking, illustrating the alignment and somatic variant-calling evaluation. **b)** Strelka2 outperforms other tools when aligned to the pangenome reference. Boxplot shows F1 score, precision, and recall (y-axis) for Somatic Sniper, Strelka2, and Mutect2 (x-axis) using pangenome alignment. Boxplots show the median (center line), interquartile range (box), and 1.5 x IQR whiskers. **c)** Strelka2 outperforms Somatic Sniper and Mutect2 when aligned to the pangenome reference compared with the linear reference. Delta F1-score, precision and recall of pangenome versus linear (y-axis) for each of the three callers (x-axis). **d)** Strelka2 shows minimal performance loss relative to consensus when using the pangenome instead of the linear reference. **e)** Comparison of pangenome and linear references when calling SNVs using consensus approach.

To assess if improved alignments resulted in improved somatic SNV detection, we next ran three somatic SNV detection algorithms, Strelka2^9^, Mutect2^10^ and Somatic Sniper^11^, on both the linear and pangenome aligned tumours (**Figure 1a; Supplementary Figure 1a**). Because current somatic mutation tools cannot be run directly on the variation graph aligned data, reads aligned to the pangenome were first projected down to the linear reference. This projection served to maintain the improved alignment from the pangenome while ensuring compatibility with existing somatic mutation detection tools. Next, we leveraged the set of SNVs detected by the TCGA MC3 for the same 30 tumours as a gold standard. Informed by crowd-sourced benchmarking efforts^12^, the MC3 took a comprehensive approach to somatic SNV detection, identifying a high confidence set of mutations based on the consensus of five distinct algorithms^13^. Such consensus approaches are time and computationally expensive. We hypothesized that the increased representation of diverse haplotypes in the pangenome would improve the precision of somatic mutation calling from a single algorithm mitigating the need for multiple algorithms and speeding up mutation detection.

### The human pangenome increases the precision of somatic SNV detection

For each somatic mutation detection tool and reference genome (*i.e*. linear vs pangenome), we assessed the performance of somatic SNV detection against the MC3 gold standard evaluating based on precision, recall and F1-score. Overall, Strelka2 run with the pangenome reference demonstrated superior F1-score compared to Mutect2 and Somatic Sniper and compared to all three tools run with the linear reference (**Figure 1b; Supplementary 3a**). The improvement in Strelka2’s F1-score was attributed to increased precision without sacrificing recall. Across all three callers, Strelka2 demonstrated the largest improvement with the pangenome compared to the linear reference (**Figure 1c**).

We next assessed if the majority vote across the three algorithms, referred to as the consensus approach, would also significantly improve precision and F1-score when using the pangenome (**Figure 1a**). Notably, the consensus approach, when aligned to the pangenome, did not significantly outperform Strelka2 with the pangenome (Effect Size (ES) = 0.645; P = 0.72; Mann-Whitney Rank Sum Test; **Figure 1d**; **Supplementary Figure 3b-c**). Indeed, the improvement in F1-score from the consensus approach was significantly smaller when aligned to the pangenome vs the linear reference (ES = -7.06; P = 2.51x10^-5^; Mann-Whitney Rank Sum Test; **Figure 1d**). On average, there was no improvement of the consensus approach leveraging the pangenome compared to the linear reference (**Figure 1e**; **Supplementary Figure 3b-c**). These data indicate that the improvement in precision conferred by the pangenome reference reduces the need for computationally expensive consensus approaches.

### Improved accuracy preferentially observed in individuals of East Asian ancestry

Next, we investigated if somatic SNV detection leveraging the pangenome differed across ancestry populations. F1-score, precision and recall were highest amongst individuals of European ancestry, followed by African and East Asian ancestry (**Figure 2a**). However, the improvement in F1-score when aligned to the pangenome was highest in individuals of East Asian ancestry (**Figure 2b**; **Supplementary Figure 4a**). Indeed, we observed on average a 20% increase in F1-score in individuals of East Asian ancestry. In contrast, improvements in individuals of European ancestry were marginal. These findings were also observed from Mutect2 and Somatic Sniper (**Supplementary Figure 4b-e**) and were replicated when considering ancestry along a continuous spectrum (**Supplementary Figure 4f-h**). These findings were also robust to tumour purity, somatic mutation burden or sequencing coverage (**Supplementary Figure 5**).

**Figure 2.**
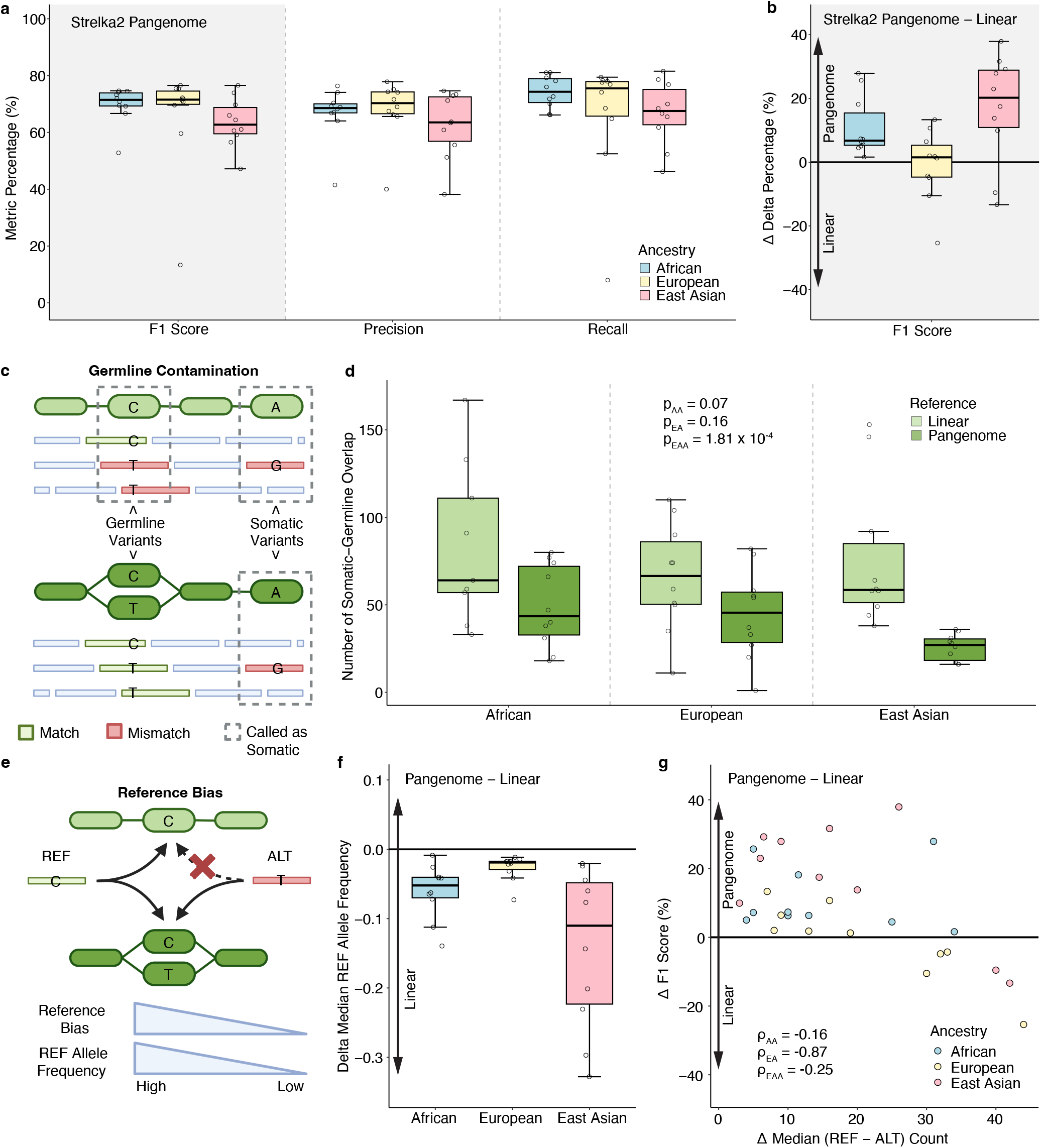
Individuals of East Asian ancestry benefit the most from the human pangenome reference. **a)** F1 score, precision, and recall for Strelka2 across ancestries using pangenome alignment. Boxplots show the median (center line), interquartile range (box), and 1.5×IQR whiskers. **b)** Individuals of Asian ancestry exhibit the largest F1 score increase with pangenome versus linear reference, with smaller increases observed for other ancestries. Boxplots show the difference in Strelka2 F1 scores for pangenome versus linear references (y-axis) across ancestries (x-axis). **c)** Schematic illustrating germline contamination and somatic variant calling in the context of pangenome and linear references. **d)** All ancestries show significantly fewer somatic-germline overlaps when aligned to the pangenome compared with the linear reference. Boxplots show somatic-germline overlap counts (y-axis) across ancestries (x-axis) for both references. For visualization purposes, one outlier was removed and the full cohort is shown in **Supplementary Figure 6a. e)** Schematic illustrating reference bias in linear genomes and how the pangenome can mitigate reference-biased allele representation. **f)** Delta median reference-allele frequency across ancestries, showing consistently lower reference-allele frequencies under the pangenome reference, indicative of less reference bias. **g)** Scatterplot of delta F1 score (y-axis) and delta median (ref - alt) count (x-axis), where larger (ref - alt) biases correspond to stronger reference bias. Lower reference bias is associated with higher F1 scores when using the pangenome relative to the linear reference.

### The human pangenome reduces germline contamination and reference bias

To determine the features driving the improved precision from the pangenome, we evaluated two known limitations with the linear reference genome: germline contamination and reference bias. Germline contamination in somatic mutation detection refers to the mislabeling of germline variants as somatic mutations (**Figure 2c**). This can occur if the germline variant is not sufficiently represented in the sequencing of the non-malignant tissue used as the matched normal. To evaluate germline contamination, we calculated the number of SNVs detected that overlapped reported sites of germline variation in gnomAD^14^. We found significantly more overlaps with reported germline sites amongst somatic SNV calls leveraging the linear reference compared to the pangenome reference (FC = 1.84; P = 1.82x10^-6^; Wilcoxon Signed-Rank Test; **Figure 2d**; **Supplementary Figure 6a-b**). More, the reduction in germline contamination was greater in individuals of East Asian ancestry compared to European ancestry (FC = 1.47; P = 0.11; Wilcoxon Signed-Rank Test; **Figure 2d; Supplementary Figure 6c-d**) helping to explain the increased somatic SNV detection accuracy in this population. Taken together, these data implicate a reduction of germline contamination as one reason why somatic SNV detection leveraging the pangenome outperforms the linear reference genome.

Reference bias occurs when reads containing alleles that do not match the reference genome fail to map correctly. This leads to a decreased representation of alternative alleles in the aligned genome. To quantify reference bias, at each site we calculated the difference in the number of reads mapping to the reference allele vs the alternate allele – the greater the delta read count (n_Ref_ – n_Alt_) the greater the reference bias (**Figure 2e**). Overall, compared to the linear genome, the pangenome reduced reference bias, particularly in individuals of East Asian descent (**Figure 2f**). Indeed, the reduction in reference bias correlated with the observed improvement in accuracy in these tumours (**Figure 2g**). To ensure somatic variant clonality was not driving these associations, we replicated these findings considering germline variants within +/- 75bp of the somatic SNVs (**Supplementary Figure 7**). Thus, the pangenome improves somatic SNV detection by reducing reference bias, particularly in individuals of non-European descent.

### Evaluating the consequences of SNV detection with the pangenome

To evaluate the consequences of SNV detection with the pangenome, we quantified MC3-exclusive (*i.e*. found by the MC3 but not the pangenome) and pangenome-exclusive SNVs (*i.e*. found by the pangenome and not MC3) in cancer driver genes, considering bladder-specific and pan-cancer driver genes^5^. For context, we identified 579 shared SNVs (*i.e*. found by both MC3 and pangenome) across all tumours in 33 bladder and 95 pan-cancer driver genes, of which 486 were predicted to have high or moderate impact. In contrast, we found 68 MC3-exclusive and 11 pangenome-exclusive SNVs in 17 bladder cancer and 34 pan-cancer driver genes across all tumours (**Supplementary Figure 8; Supplementary Table 2**). Considering only SNVs predicted to be moderate or high impact, we found 31 MC3-exclusive and 3 pangenome-exclusive SNVs in 9 bladder cancer and 18 pan-cancer driver genes (**Figure 3a-b**). Thus, the SNVs that are not concordant between the MC3 and the pangenome represent only 6.5% of the high or moderate impact SNVs identified in cancer driver genes.

**Figure 3.**
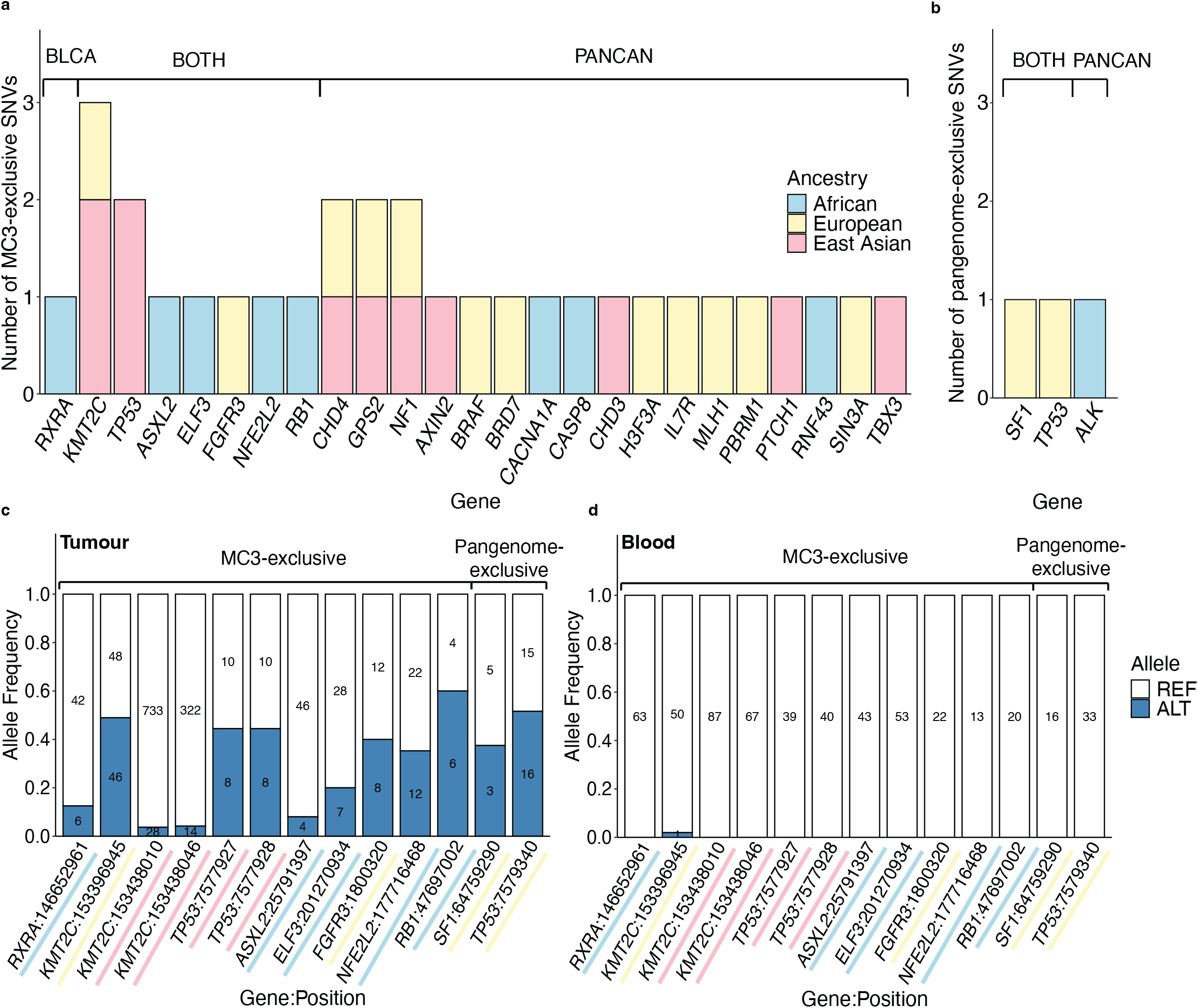
Consequences of SNV detection leveraging the pangenome. **a)** Number of predicted high or moderate-impact MC3-exclusive SNVs in bladder or pan-cancer driver genes coloured by genetic ancestry. **b)** Number of predicted high or moderate-impact pangenome-exclusive SNVs in bladder or pan-cancer driver genes coloured by genetic ancestry. **c-d)** Frequency of reads mapping to the alternative allele (y-axis) at each SNV (x-axis) in high or moderate-impact variants in bladder cancer driver genes in tumour **(c)** and blood **(d)** sequencing. Numbers within the bars indicate the read counts supporting the alternative (blue) or reference (white) alleles. Coloured underline beneath x-axis labels indicates the ancestry population that the SNV was found in.

While the MC3 represents a rigorous, expert curated list of SNVs, it may still not reflect the ground truth. Therefore, to determine if MC3-exclusive SNVs represent true somatic SNVs, we examined read-level evidence. Focusing on moderate and high-impact SNVs in bladder cancer driver genes, we quantified the number of reads that support the reference and alternate allele at each site in both the tumour and blood sequencing aligned to the pangenome. A true somatic SNV in the tumour is expected to show multiple reads supporting the alternative allele in the tumour sequencing and zero reads supporting the alternative allele in the blood sequencing. This held true for the MC3-exclusive SNVs (**Figure 3c-d**), with the exception of a SNV in *KMT2C* (chr7:153396945) which showed only a single read supporting the alternative allele in the blood sequencing and likely a sequencing artifact. Half of the MC3-exclusive SNVs had alternative allele frequencies ranging from 0.4-0.6 suggesting they are true clonal somatic SNVs, while the other half had allele frequencies ranging 0.04-0.38 aligned with subclonal somatic SNVs. Two *KMT2C* variants (chr7:153396945 and chr7:153396945) in two separate individuals had low alternative allele frequencies (0.04) but still had 14-28 reads supporting the alternative allele (**Supplementary Table 2**). In total, these loci had 336-761 reads mapped. Thus, these two SNVs are likely true low frequency subclonal SNVs occurring within a region of the genome that has been amplified. Taken together, these data indicate that the improved precision with the pangenome can lead to a small subset (<4%) of high or moderate impact SNVs in bladder cancer driver genes being overlooked.

Importantly, not all MC3-exclusive SNVs represent limitations of the pangenome. There were two variants in *TP53* found in a single tumour by the MC3 and not the pangenome (TCGA-CF-A1HS; **Figure 3c-d**). These two variants were adjacent representing a dinucleotide variant (CG>TT), rather than an SNV. This variant was not detected by Strelka2 because Strelka2 does not support dinucleotide variant detection. Given that Mutect2 does, we confirmed that Mutect2 with pangenome alignment correctly identified the CG>TT dinucleotide variant in TCGA-CF-A1HS. Thus, the *TP53* variant does not reflect an error with the pangenome alignment but rather highlights the importance of selecting the appropriate algorithm for the mutation type of interest.

In contrast to the MC3-exclusive SNVs, the pangenome-exclusive SNVs in bladder cancer driver genes provided evidence that the pangenome recovers variants absent from MC3. Quantifying allele frequencies for high or moderate pangenome-exclusive variants in bladder cancer driver genes found all SNVs showed ample read support for the alternative allele in the tumour (allele frequency = 0.38-0.51), and no alternative allele read support in the normal (**Figure 3c-d; Supplementary Table 2**). These data support both variants as likely true SNVs that were missed by the MC3 and suggest the pangenome may recover true somatic SNVs absent from curated callsets.

### Improved precision leveraging the pangenome generalizes to lung adenocarcinoma

Finally, to evaluate if the improvements we observed with the human pangenome extended beyond bladder cancer, we benchmarked somatic SNV detection in 29 lung adenocarcinoma tumours from TCGA, with equal representation of donors of European, African and East Asian ancestry, though only nine donors of East Asian ancestry were available (**Supplementary Figure 9**). Alignment to the pangenome showed no significant difference in the number of aligned reads, however, the proportion of properly paired reads was significantly increased, in concordance with our observations in bladder tumours (FC = 1.02; P = 1.23x10^-12^; Wilcoxon Signed-Rank Test; **Supplementary Figure 10**). Next, we investigated if somatic SNV detection using Strelka2 and the pangenome outperformed the linear reference in lung tumours (**Figure 4a**). There were three tumours with F1-score < 20% when aligned to both the linear and pangenome references. These tumours had fewer than 10 SNVs detected by MC3 which led to low precision with both alignments. Overall, Strelka2 with the pangenome showed improved F1-score compared to the linear reference (**Figure 4b-c; Supplementary Figure 11a**) primarily driven by an increase in precision (**Figure 4c**), similar to bladder tumours. Improvements were disproportionally higher for individuals of East Asian ancestry (**Figure 4d-e; Supplementary Figure 11b**), while improvements in individuals of European ancestry were marginal (**Figure 4e**). These findings demonstrate that broad adoption of the human pangenome has the potential to meaningfully reduce ancestry-related disparities in somatic mutation detection, with the greatest gains expected for individuals currently underrepresented in the linear reference genome.

**Figure 4.**
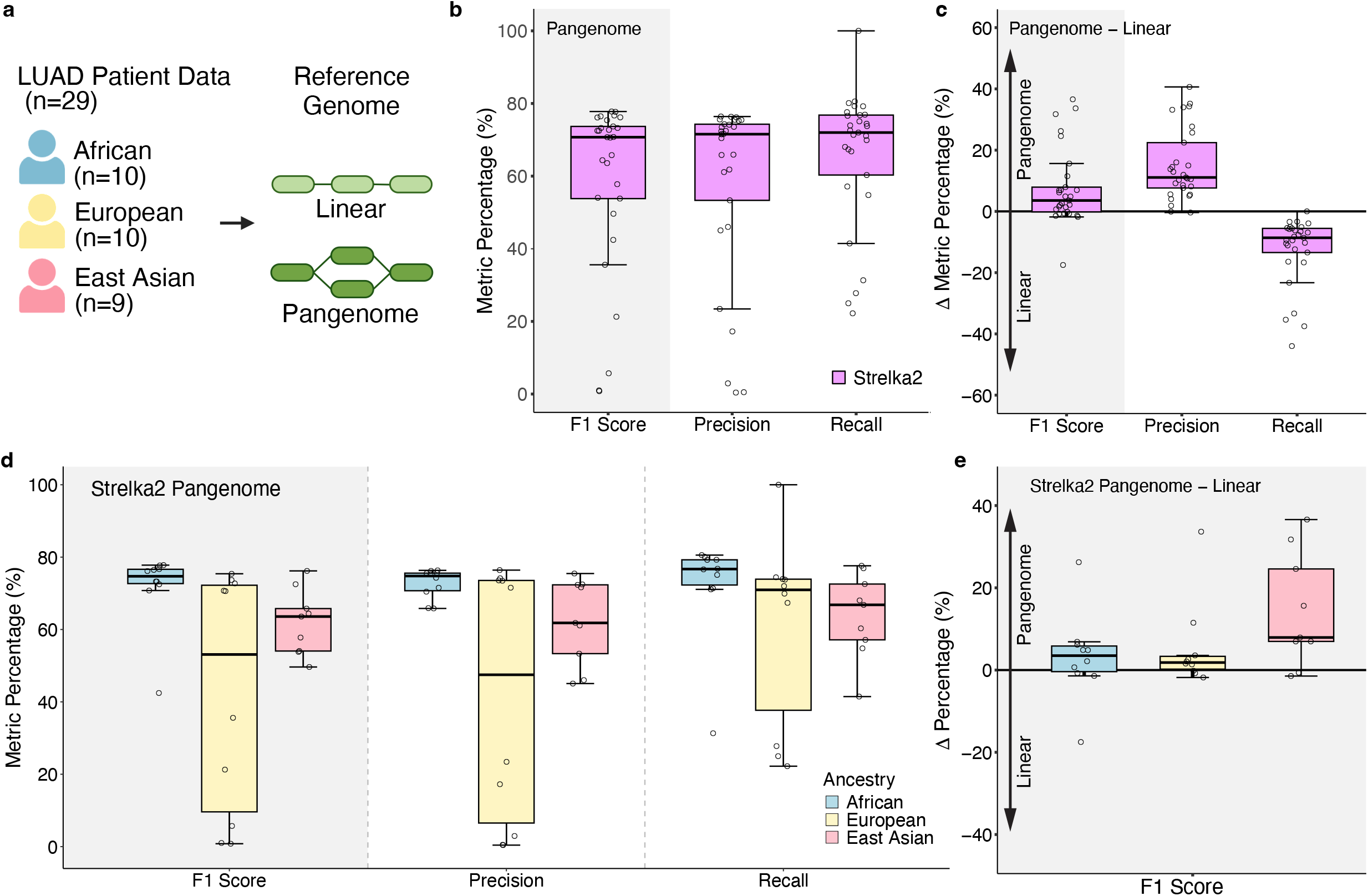
Increased precision leveraging the pangenome generalizes to lung adenocarcinoma. **a)** Schematic overview of benchmarking in lung adenocarcinoma tumours. **b)** Boxplot shows F1 score, precision, and recall (y-axis) for Strelka2 (x-axis) using pangenome alignment in lung tumours. Boxplots show the median (center line), interquartile range (box), and 1.5 x IQR whiskers. **c)** Strelka2 when aligned to the pangenome reference outperforms the linear reference. Delta F1-score, precision and recall of pangenome versus linear (y-axis). **d)** F1 score, precision, and recall for Strelka2 across ancestries using pangenome alignment in lung tumours. **e)** Boxplots show the difference in Strelka2 F1 scores for pangenome versus linear references (y-axis) across ancestries (x-axis) in lung tumours.

## DISCUSSION

There has been a strong push to increase representation in genomics with a focus on increasing ancestry representation in genomics cohorts. Less attention has been given to representation in the methods used to analyze genomic data. Here, we demonstrate that leveraging the human pangenome reference in somatic SNV detection can help to mitigate ancestry-related biases in somatic mutation detection. Specifically, we show that the human pangenome reference improves the precision of somatic mutation detection, particularly in individuals of East Asian ancestry, who are least represented by the linear reference^1^. Indeed, we observed on average a 20% improvement in F1-score in individuals of East Asian ancestry when leveraging the human pangenome compared to the linear genome. In contrast, observed improvements for individuals of European ancestry were more modest. These differences in performance across ancestries was observed in both bladder carcinoma and lung adenocarcinoma. Overall, the increase in precision can be attributed to both a reduced germline contamination along with reduced reference bias.

Further, we found somatic SNV detection with Strelka2 leveraging the human pangenome did not outperform the consensus approach. Previous international crowd-sourced benchmarking studies of somatic SNV detection leveraging the linear genome found that a “wisdom-of-the-crowds” approach substantially improved precision^12^. However, these approaches require running multiple tools and can be both time and computationally expensive, particularly as cancer genomics cohorts now consist of thousands of tumour samples. Here, we demonstrate that with the pangenome, these computationally expensive consensus approaches may no longer be required. Instead, the pangenome, along with Strelka2, improves precision comparability.

Notably, we observed superior performance using Strelka2 compared to Mutect2. Differences in performance may reflect differences in the external resources used by each caller rather than fundamental differences between the tools. Specifically, Mutect2 incorporates germline population allele frequency priors derived from gnomAD and a panel of normals, both of which were generated using linear reference genome alignments. In contrast, Strelka2 relies primarily on differences in allele frequencies between the tumour and normal samples, reducing its dependence on external population priors that may be mismatched to the pangenome reference. These results suggest that when adopting a new reference genome, careful consideration should be given to whether the accessory resources are appropriate for the new alignment context.

Importantly, the observed improvements to somatic mutation detection accuracy were observed when only considering the coding genome which represents 2% of the most well-characterized regions of the genome. We anticipate larger improvements in non-coding regions of the genome and when considering larger more complex alterations, such as structural variants. Indeed, the human pangenome consortium found 104% increase in structural variant detection compared to a 34% improvement in single nucleotide polymorphism detection^2^. Thus, we anticipate the improvements found here likely serve as a lower bound.

Finally, a current limitation is that somatic mutation callers cannot operate directly on variation graph alignments, requiring reads to be projected back to the linear reference prior to variant calling. This projection preserves the benefits of pangenome alignment but introduces mismatches at sites where a read more closely resembles an alternative haplotype than the linear reference. As fully pangenome-aware variant callers are developed, we anticipate further improvements in somatic mutation detection. In the interim, our findings provide practical guidance for minimising ancestry-related biases in somatic variant calling pipelines.

## CONTRIBUTIONS

Performed statistical and bioinformatic analyses: CP, FA, TH, EA, AB, ES

Wrote first draft of the manuscript: KEH

Edited manuscript: CP, FA, TH, EA, AB, ES, KEH

Initiated the project: FA, KEH

Supervised research: KEH

Approved the manuscript: All authors

## DATA AVAILABILITY

Whole exome sequencing for the tumour and matched blood sequencing for 30 bladder carcinoma and 29 lung adenocarcinoma tumours evaluated here can be found on GDC (https://portal.gdc.cancer.gov/).

## CODE AVAILABILITY

Code will be made available on GitHub. A Docker image of the pangenome and Strelka2 workflow will be made available on DockerHub.

## ACKNOWLEDGEMENTS

This research was supported by NSERC through a Discovery Grant (RGPIN-2025-05991 and DGECR-2025-00288) as well as a Canada Research Chair Award (CRC-2024-00022) to KEH. The researchers would also like to acknowledge the research was made possible by patients and families that participated in the TCGA PanCancer Atlas.

