## Supplementary Materials for "The human pangenome reference reduces ancestry-related biases in somatic mutation detection"

Supplementary Figure 1

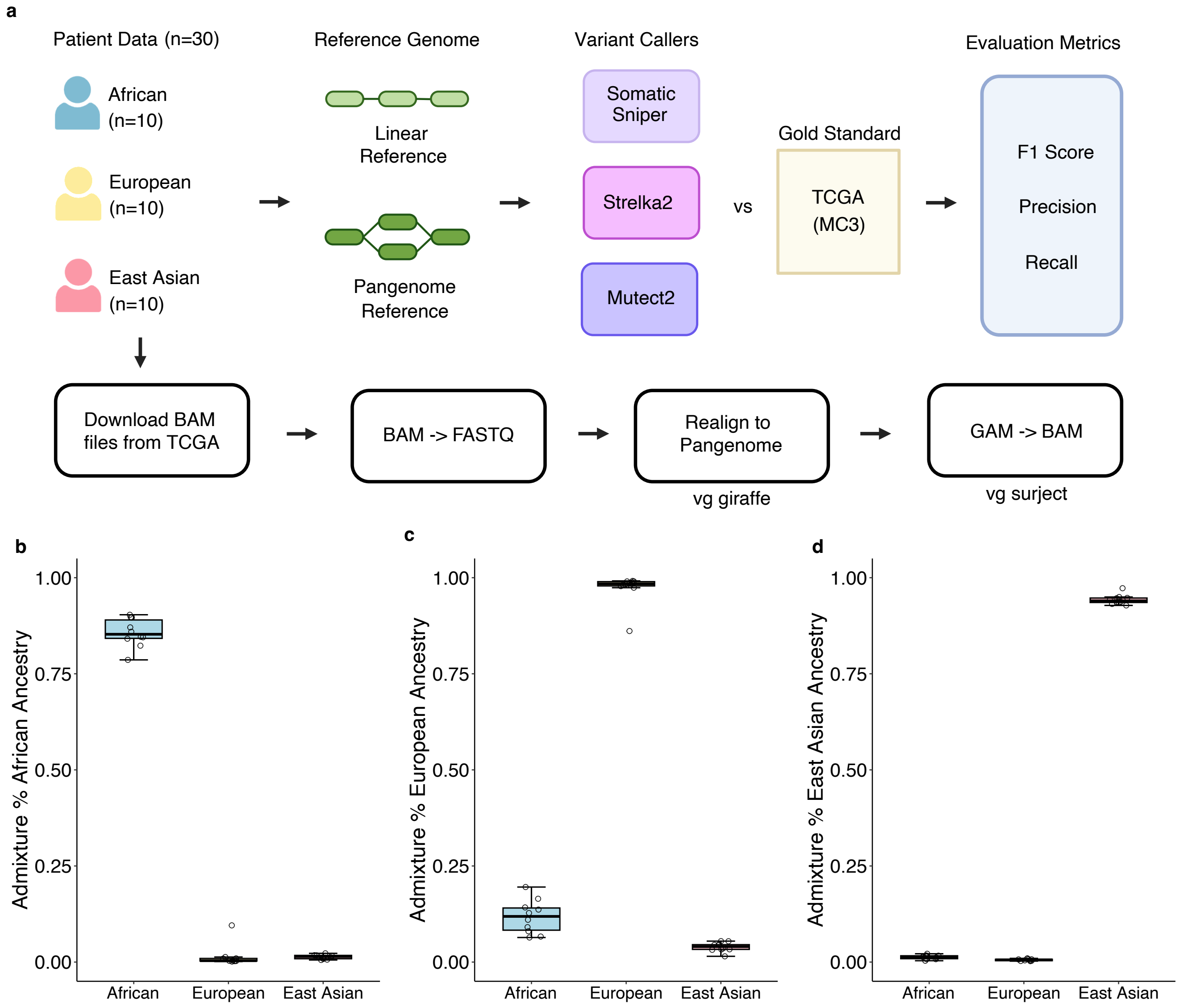

**Supplementary Figure 1. Cohort and benchmark overview**

**a)** Schematic overview of the alignment, variant calling, and evaluation steps. **b)** African ancestry samples show highest admixture percentage in African ancestry. **c)** European ancestry samples show highest admixture percentage in European ancestry. **d)** East Asian ancestry samples show highest admixture percentage in East Asian ancestry. Boxplots show the median (center line), interquartile range (box), and 1.5 x IQR whiskers.

Supplementary Figure 2

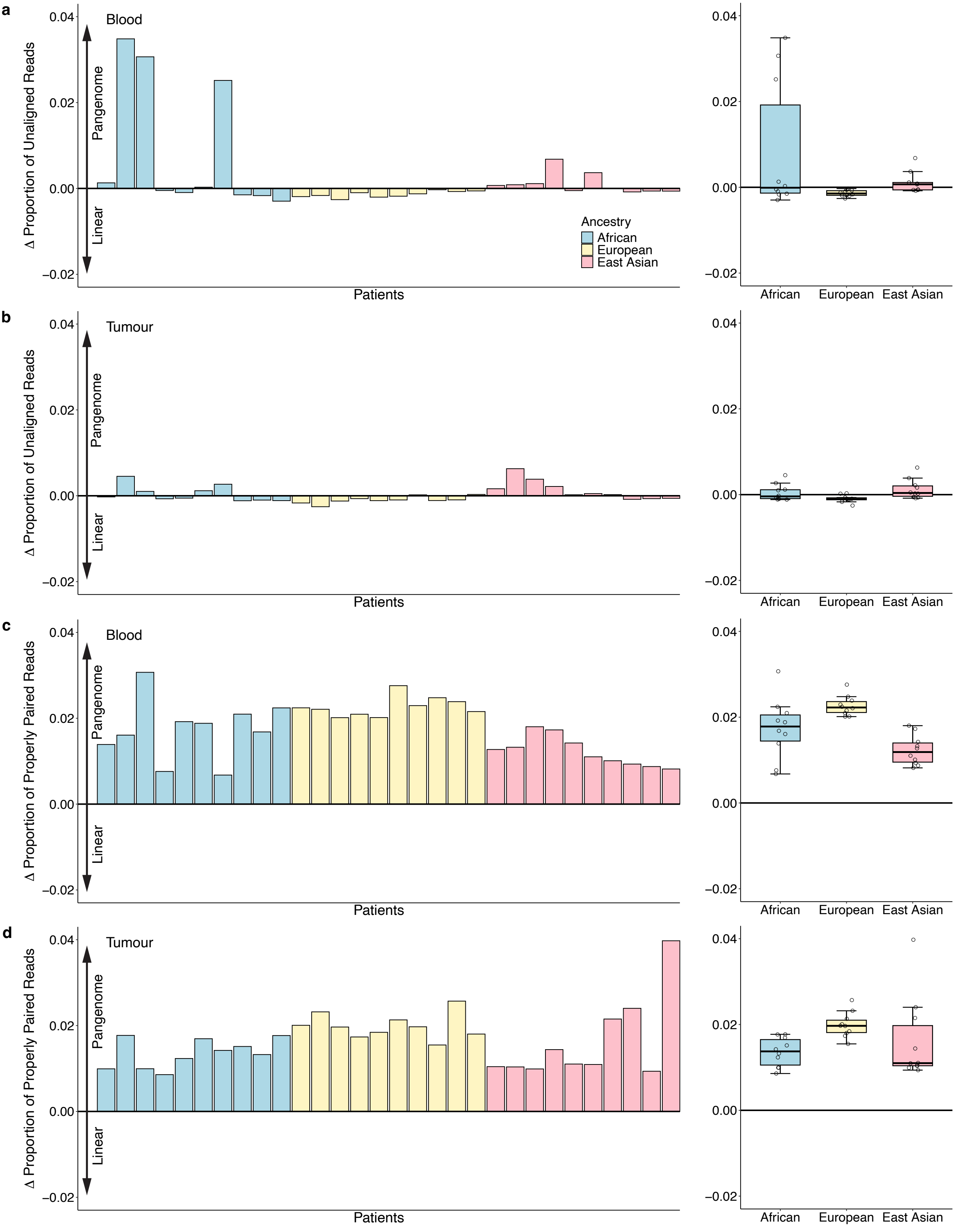

**Supplementary Figure 2. The human pangenome improves DNA alignment accuracy**

**a-b)** Change in the proportion of unaligned reads when mapping blood **(a)** and tumour **(b)** samples to the pangenome compared with the linear reference, shown per patient (left) and summarized by ancestry (right). Bars represent individual patients coloured by ancestry; boxplots show distributions for African, European, and East Asian groups. **c-d)** Change in the proportion of properly paired reads in blood **(c)** and tumour **(d)** samples when aligned to the pangenome relative to the linear reference. Positive values indicate an increase in properly paired reads under the pangenome reference. Boxplots (right) summarize the distribution of differences across ancestries. Across both blood and tumour datasets, African ancestry samples exhibit the largest shifts in alignment metrics when using the pangenome.

Supplementary Figure 3

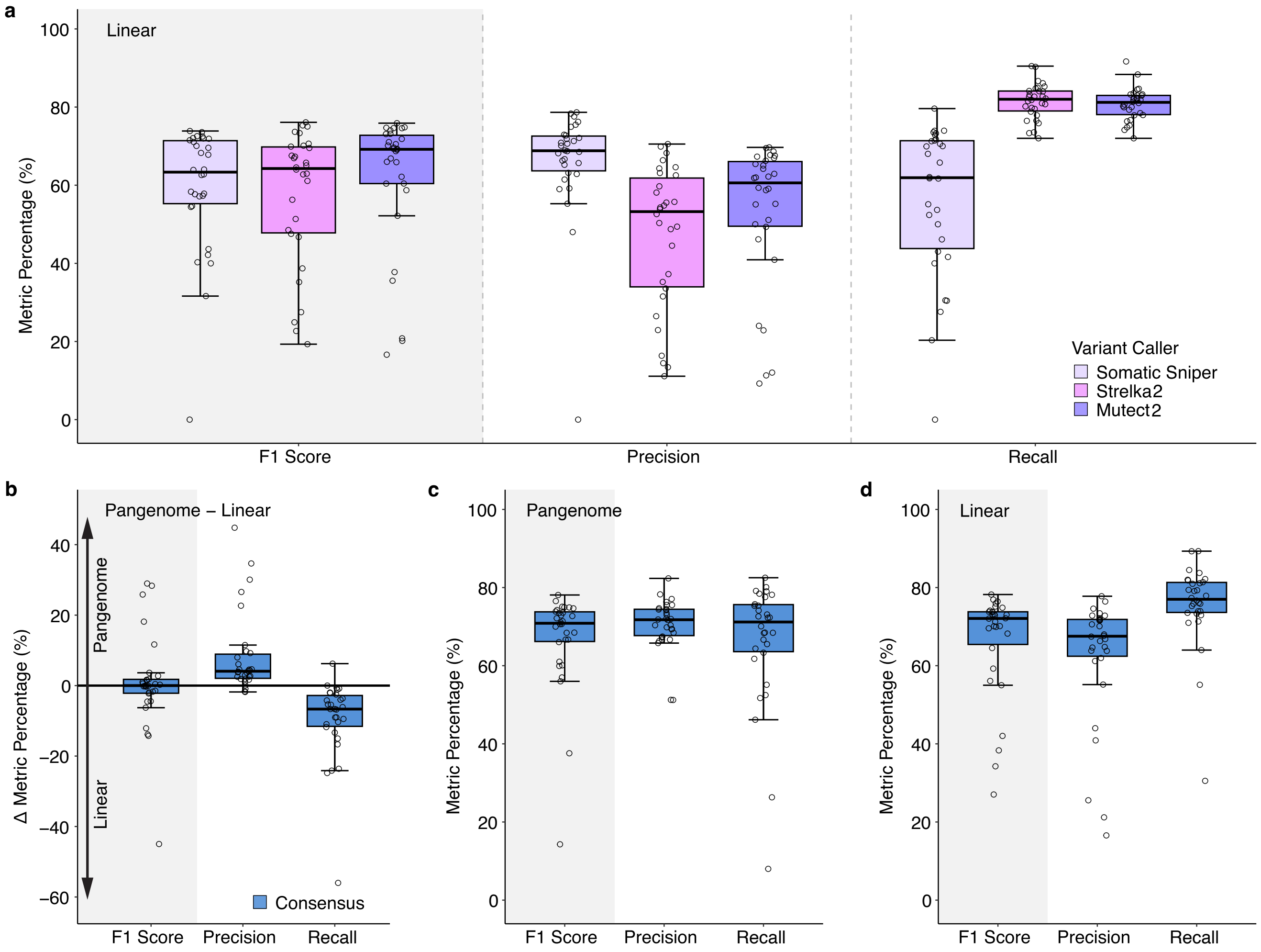

**Supplementary Figure 3. The human pangenome improves the precision of somatic mutation detection**

**a)** Boxplots show F1 score, precision, and recall for different variant caller samples when aligned to the linear reference. **b-c)** F1 score, precision, and recall for consensus variant caller when aligned to the **(b)** pangenome reference and **(c)** linear reference.

Supplementary Figure 4

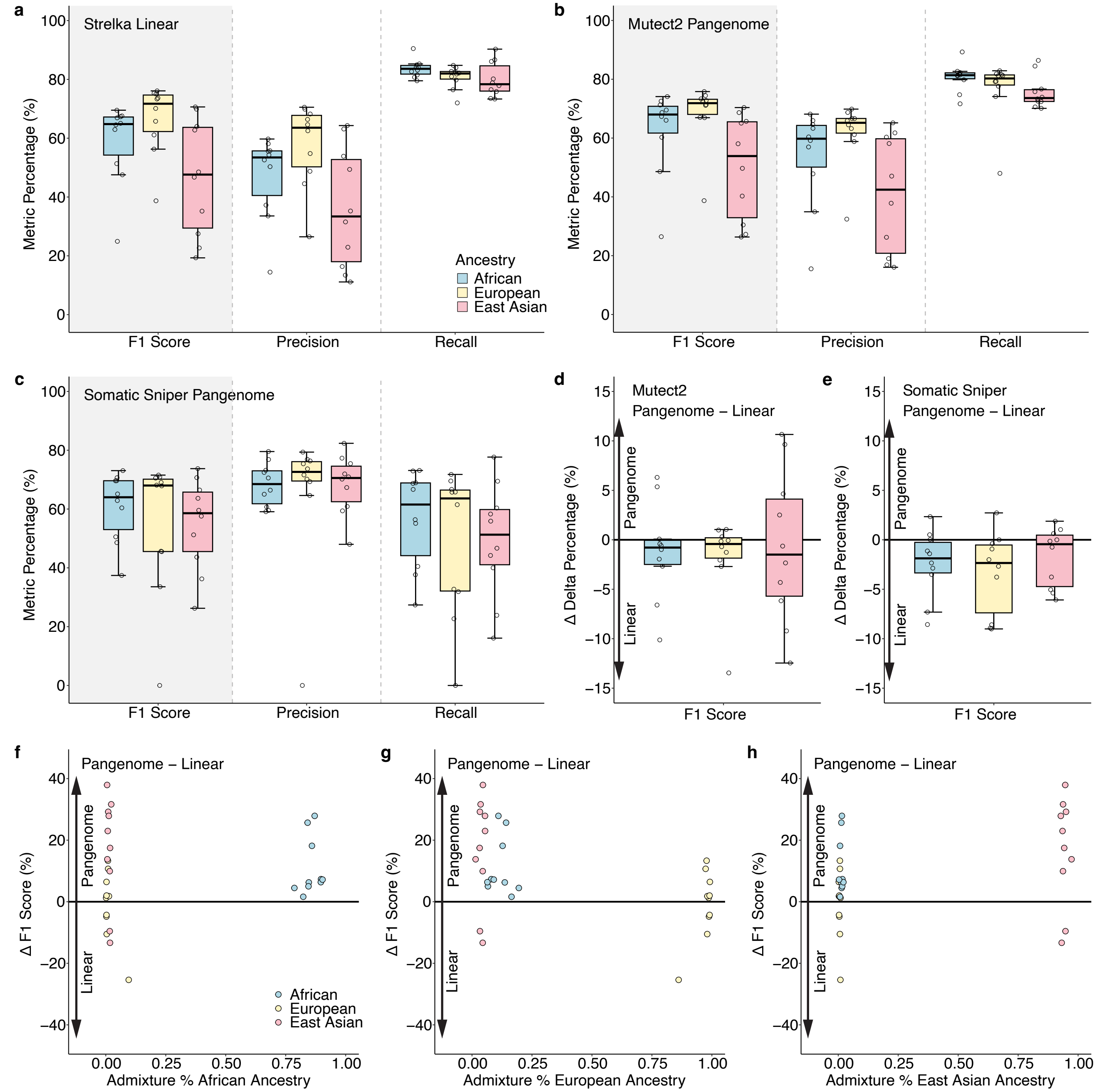

**Supplementary Figure 4. Individuals of East Asian ancestry benefit the most from the human pangenome reference**

**a-c)** Performance of different variant callers stratified by ancestry. Boxplots show F1 score, precision, and recall for African, European, and East Asian ancestry samples under **(a)** Strelka2 using the linear reference, **(b)** Mutect2 using the pangenome reference, and **(c)** Somatic Sniper using the pangenome reference. **d-e)** Change in F1 score when using the pangenome versus the linear reference for **(d)** Mutect2 and **(e)** Somatic Sniper, summarized by ancestry. Positive delta values indicate improved performance with the pangenome. **f-h)** Relationship between ancestry proportion and change in F1 score when switching from the linear to the pangenome reference. Each point represents an individual sample, coloured by reported ancestry. Panels show admixture with **(f)** African, **(g)** European, and **(h)** East Asian ancestry components.

Supplementary Figure 5

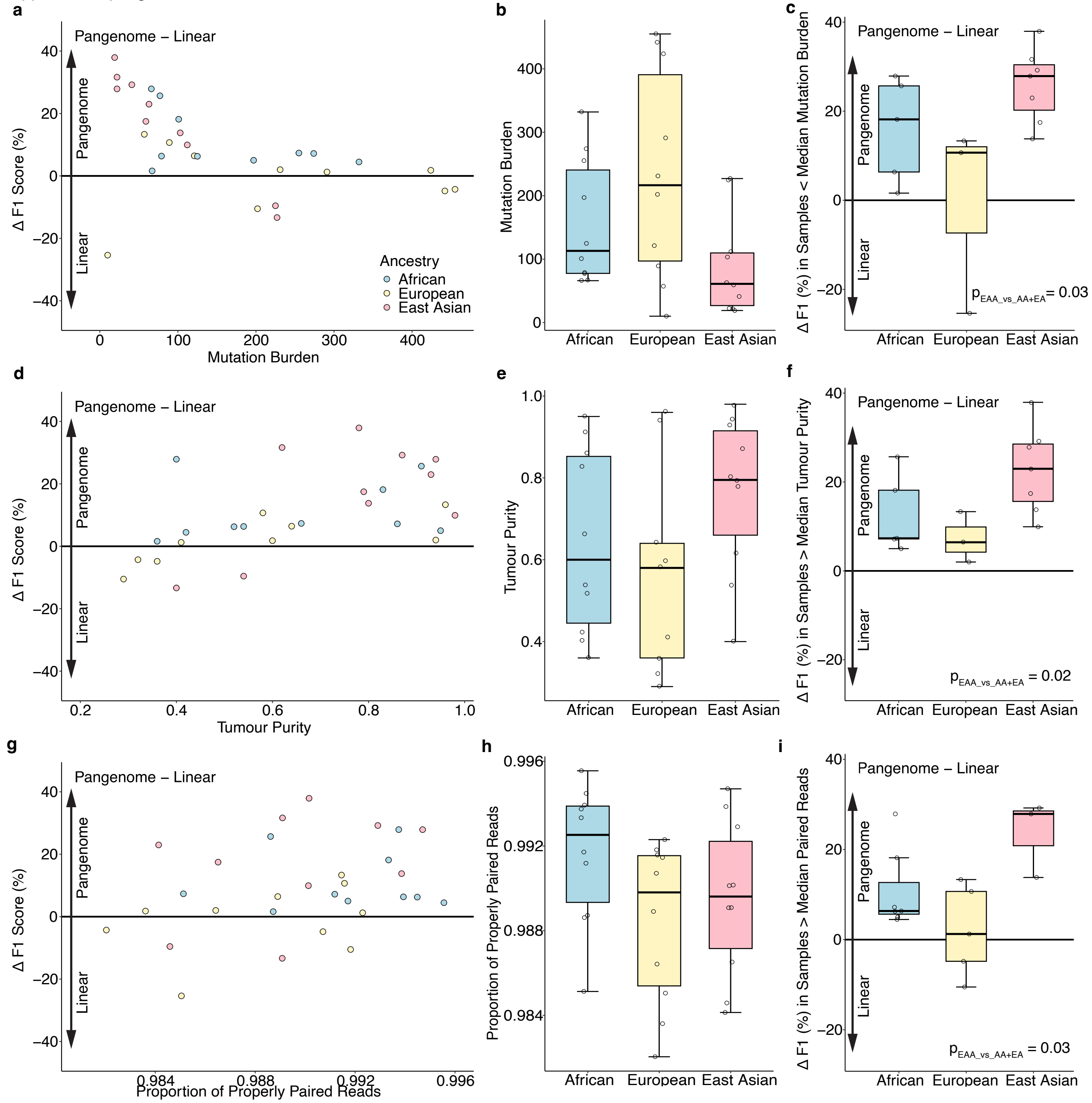

**Supplementary Figure 5. Assessment of confounders on pangenome alignment performance**

**a-c)** Delta F1 score versus mutation burden. **(a)** Scatterplot of delta F1 score against mutation burden. **(b)** Boxplots of mutation burden across ancestries. **(c)** Delta F1 scores for samples with mutation burden below the median, to adjust for differences in mutation burden in Asian samples. **d-f)** Delta F1 score versus tumour purity. **(d)** Scatterplot of delta F1 score against tumour purity. **(e)** Boxplots of purity distributions across ancestries. **(f)** Delta F1 scores for samples above the median purity, to adjust for differences in tumour purity in Asian patients. **g-i)** Delta F1 score versus proportion of properly paired reads. **(g)** Scatterplot of delta F1 score against proportion of properly paired reads. **(h)** Boxplots of paired-read proportion across ancestries. **(i)** Delta F1 scores for samples above the median paired-read proportion, to adjust for differences in paired-read proportion in Asian ancestry.

Supplementary Figure 6

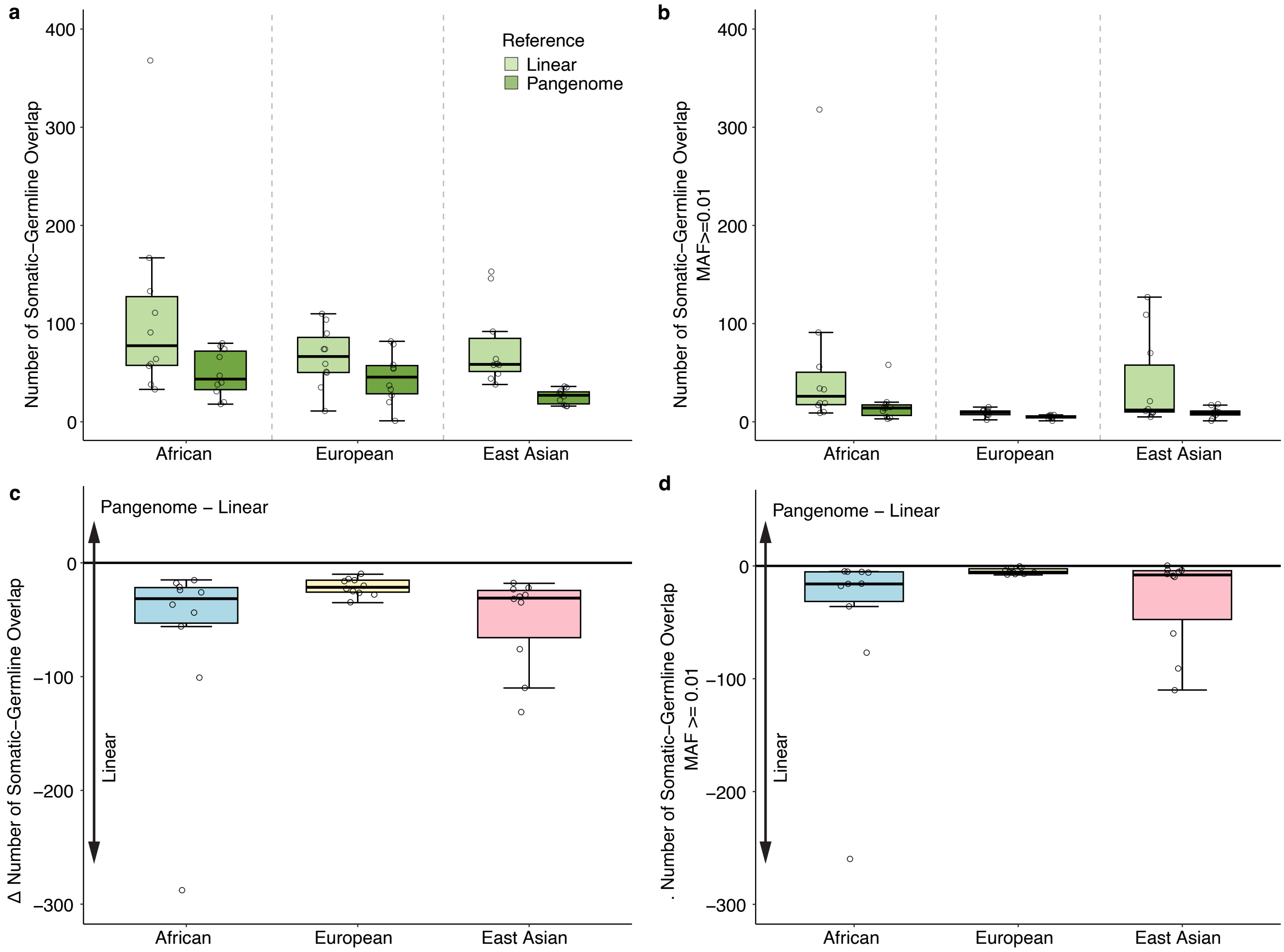

**Supplementary Figure 6. The pangenome reference reduces germline contamination**

**a-b)** Number of somatic variants overlapping germline variants (y-axis) when aligned to the linear reference or the pangenome for individuals of different ancestral groups (x-axis). **(a)** Overlap using all germline variants; **(b)** overlap restricted to germline variants with MAF $\geq 0.01$. Boxplots show the median (center line), interquartile range (box), and 1.5 x IQR whiskers. **c-d)** Change in somatic-germline overlap of pangenome minus linear for each population group. **(c)** All germline variants; **(d)** germline variants with MAF $\geq 0.01$. Negative values indicate fewer somatic variants coinciding with germline sites when using the pangenome relative to the linear reference.

Supplementary Figure 7

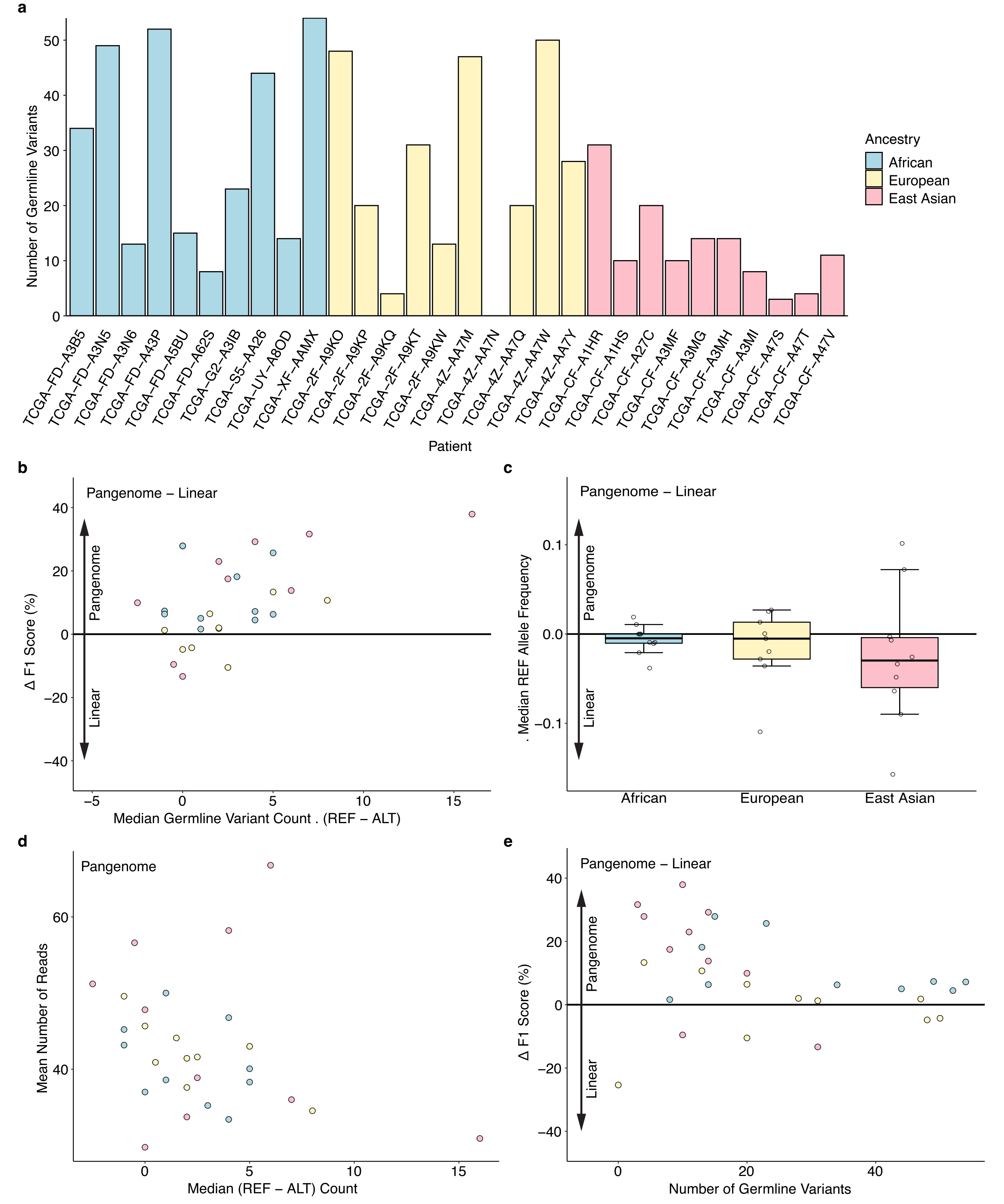

**Supplementary Figure 7. The pangenome reference reduces reference bias**

**a)** Number of germline variants within +/- 75bp of a somatic SNV per patient, stratified by ancestry. Bars represent per-sample counts for African, European, and East Asian ancestry patients. **b)** Association between change in somatic variant calling performance (y-axis) and germline variant imbalance between reference and alternate allele counts (x-axis) when using the pangenome versus the linear reference. **c)** Change in median reference allele frequency between pangenome and linear across ancestries. Boxplots show distribution of delta median REF allele frequency per ancestry. **d)** Relationship between median REF-ALT count in germline variants and mean read depth under the pangenome reference. Each point represents an individual sample colored by ancestry. **e)** Association between total germline variant count and change in F1 score of pangenome minus linear.

Supplementary Figure 8

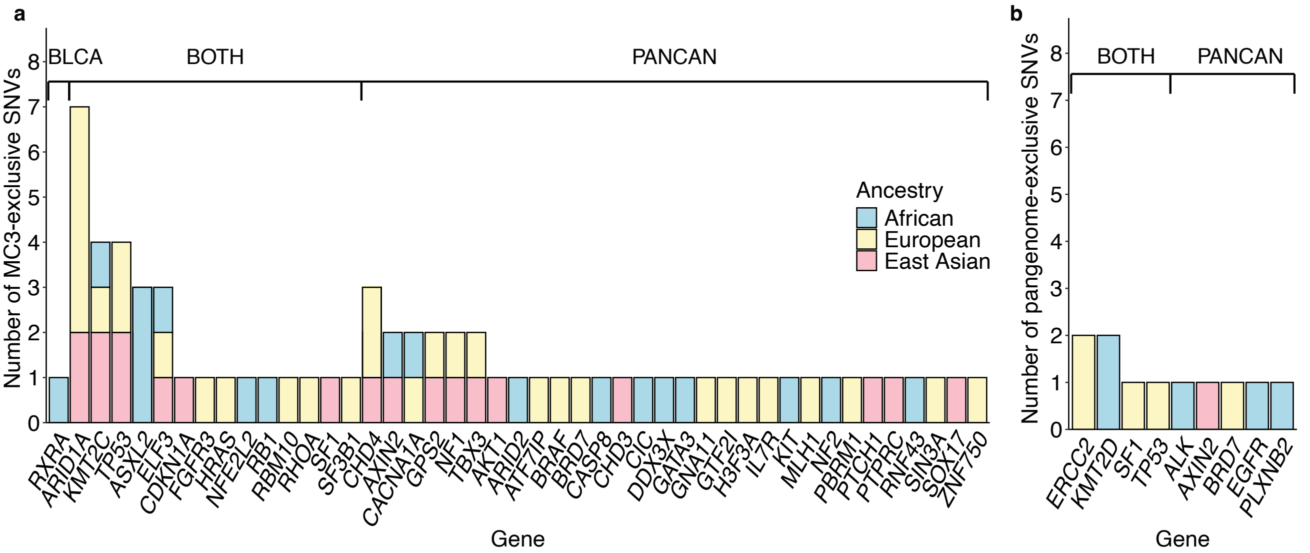

**Supplementary Figure 8. MC3-exclusive and pangenome-exclusive SNVs in cancer driver genes**

**a)** Number of MC3-exclusive SNVs in bladder or pan-cancer driver genes coloured by genetic ancestry. **b)** Number of pangenome-exclusive SNVs in bladder or pan-cancer driver genes coloured by genetic ancestry.

Supplementary Figure 9

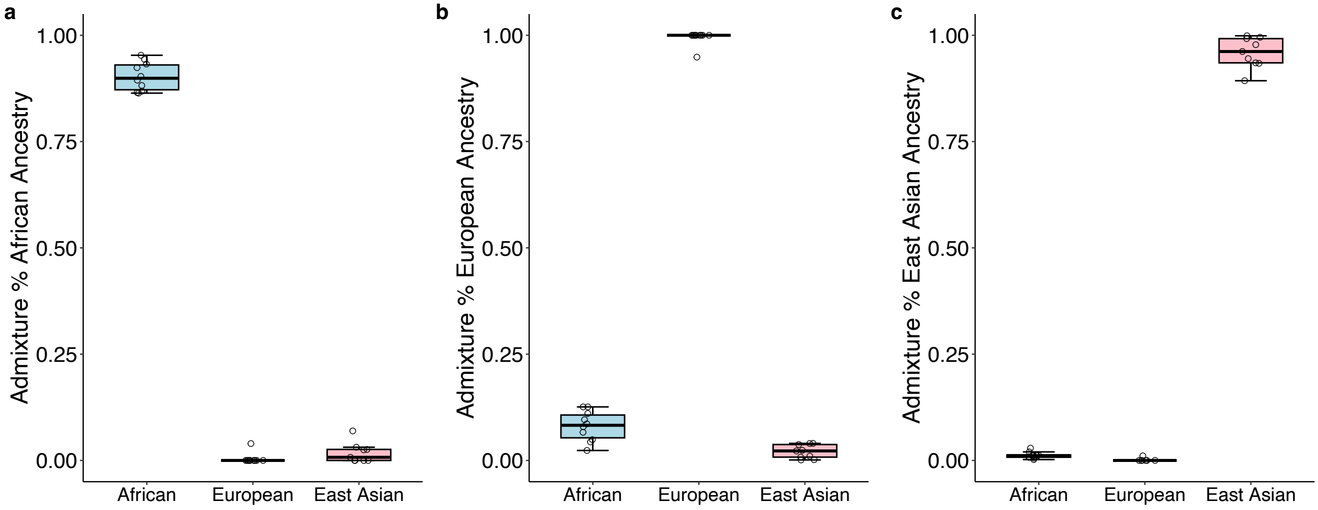

**Supplementary Figure 9. Genetic ancestry of lung adenocarcinoma tumours**

**a)** African ancestry lung tumour samples show highest admixture percentage in African ancestry. **c)** European ancestry lung tumour samples show highest admixture percentage in European ancestry. **d)** East Asian ancestry lung tumour samples show highest admixture percentage in East Asian ancestry. Boxplots show the median (center line), interquartile range (box), and 1.5 x IQR whiskers.

Supplementary Figure 10

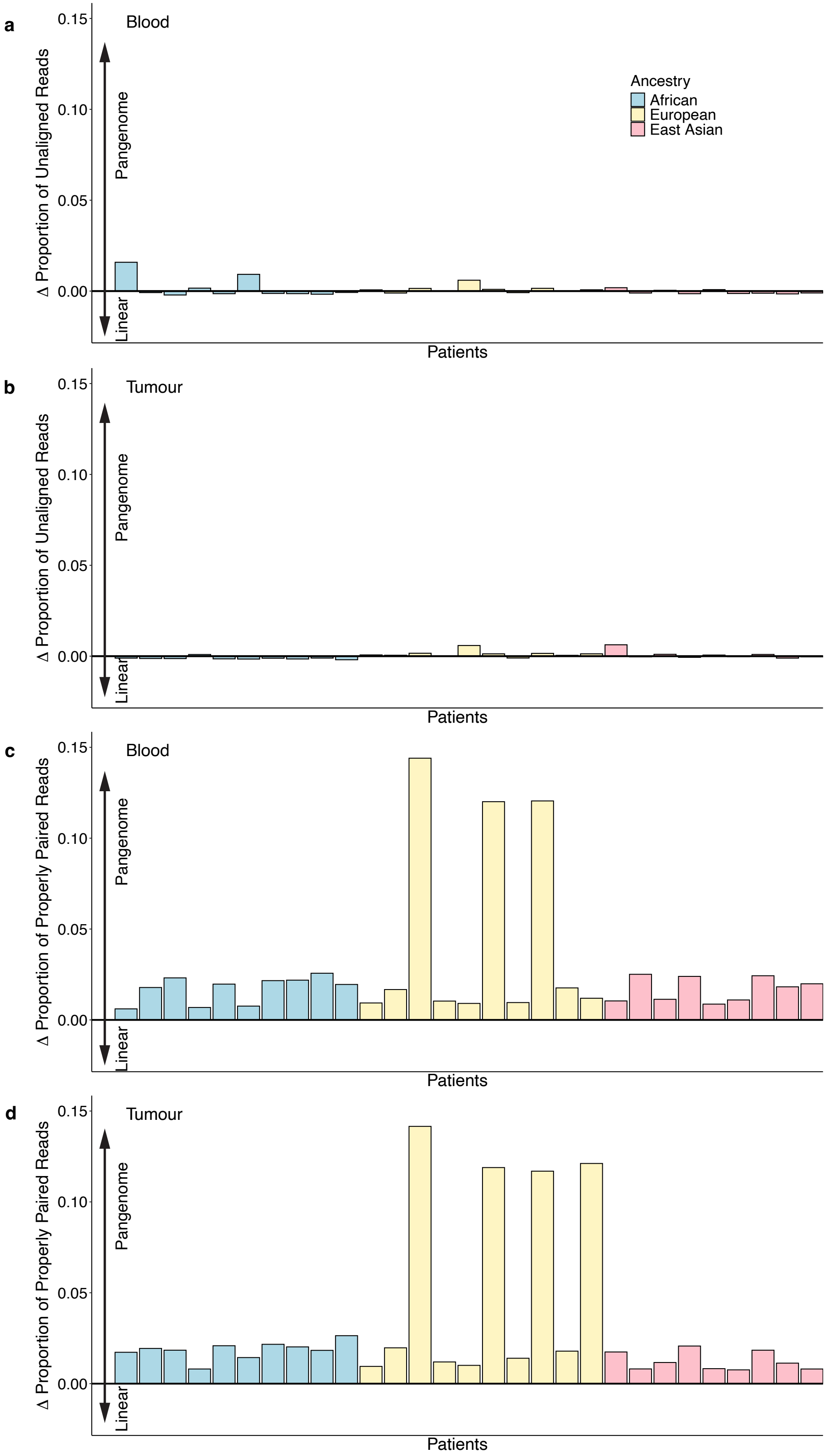

**Supplementary Figure 10. The human pangenome improves DNA alignment accuracy in lung tumours**

**a-b)** Change in the proportion of unaligned reads when mapping blood **(a)** and tumour **(b)** samples to the pangenome compared with the linear reference, shown per patient (left) and summarized by ancestry (right) in lung tumours. Bars represent individual patients coloured by ancestry. **c-d)** Change in the proportion of properly paired reads in blood **(c)** and tumour **(d)** samples when aligned to the pangenome relative to the linear reference in lung tumours. Positive values indicate an increase in properly paired reads under the pangenome reference.

Supplementary Figure 11

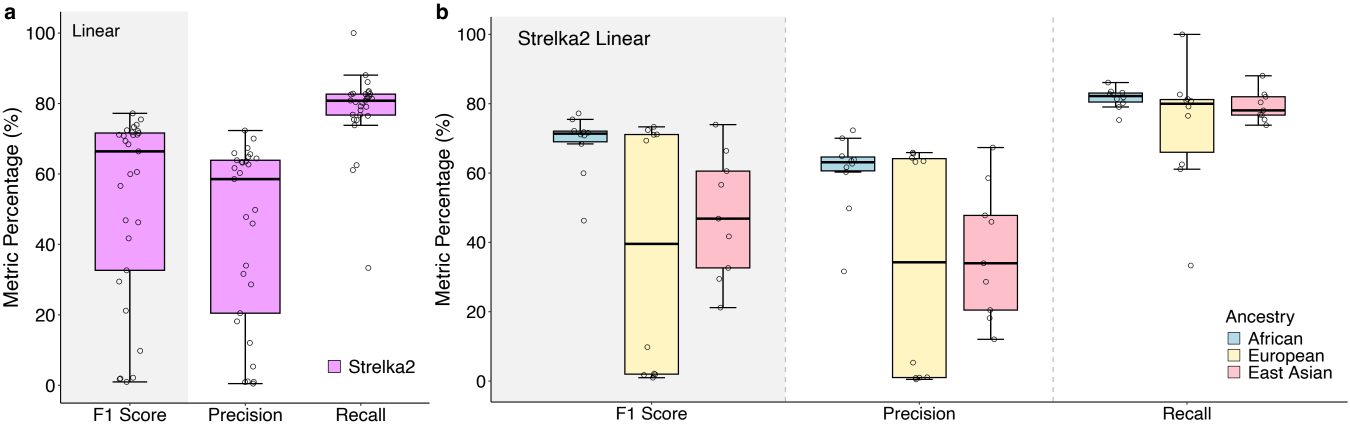

**Supplementary Figure 11. Alignment to linear reference shows reduced precision in lung adenocarcinoma**

**a)** Boxplot shows F1 score, precision, and recall (y-axis) for Strelka2 (x-axis) using linear alignment in lung tumours. Boxplots show the median (center line), interquartile range (box), and 1.5 x IQR whiskers. **b)** F1 score, precision, and recall for Strelka2 across ancestries using linear alignment in lung tumours.

### SUPPLEMENTARY TABLE LEGEND

**Supplementary Table 1. Summary of benchmarking metrics**. Benchmarking metrics across all 59 tumours ($n_{BLCA} = 30$; $n_{LUAD} = 29$) aligned to both the pangenome and linear reference genomes.

**Supplementary Table 2. MC3-exclusive and pangenome-exclusive SNVs in cancer driver genes**. Summary of high or moderate impact MC3-exclusive and pangenome-exclusive SNVs in bladder cancer driver genes, including allele frequencies in tumour and blood sequencing.

### METHODS

#### Sample Selection

We selected 30 bladder cancer patients from The Cancer Genome Atlas (TCGA) based on inferred genetic ancestry: 10 of African ancestry, 10 of East Asian ancestry and 10 of European ancestry according to Yuan *et al.*[1]. Individuals were selected based on maximum proportion of each ancestry population. Whole exome sequencing BAM files for both tumour and matched normal aligned to GRCh38 were downloaded from GDC. Reads were back-extracted from BAM files using samtools bamtofastq (v 1.16.1).

#### Alignment to Pangenome Reference

Tumour and blood whole exome sequencing was aligned to the human pangenome reference (v1.1) using *vg giraffe*[2] (v1.61.0) with the following parameters: a specified fragment length of 150 bp (-D 150) with all other parameters kept at their defaults. Because current somatic mutation tools expect alignment to a linear reference as input, we projected the variation graph aligned reads down to the T2T-CHM13 (v1.1) linear reference using *vg surject* (v1.61.0) with default parameters. A chromosome path file (-F) was added to ensure surjection mapped reads onto the correct T2T-CHM13 chromosomes. This projection converted GAM files to BAM files. We then marked duplications and ran base recalibration on the linear projected BAM file using GATK (v 4.3.0.0) MarkDuplicates and BaseQualityScoreRecalibration (BQSR), respectively. The quality of the alignment was assessed with samtools stats (v1.16.1).

#### Somatic SNV Detection

We evaluated three somatic SNV detection tools: Strelka2 (v2.9.10)[3], SomaticSniper[4] (v 1.0.5.0) and Mutect2[5] (v4.6.1.0). All three tools were run on tumour and matched normal samples for each donor. Only SNVs within the captured exome regions were considered. To identify these regions, we used panel information provided by Agilent for the SureSelect All Exon V5 library prep[6].

#### Strelka2

Strelka2[3] (v2.9.10) was run on both the linear and pangenome aligned BAM files using the somatic workflow configuration script configureStrelkaSomaticWorkflow.py and the following parameters:-callregions, -exome, and -rundir. After configuration, the workflow was executed using runWorkflow.py -m local –j 8. We only considered variants that passed the internal filters, as determined by the "PASS" annotation in the FILTER column of the resulting VCF.

#### SomaticSniper

SomaticSniper[4] (v1.0.5.0) was run on both the linear and pangenome aligned BAM files using bam-somaticsniper command and the following parameters recommended by SomaticSniper developers: -Q 40, -G, -L, -f *reference.fa*, *tumor.bam*, *normal.bam*, and -F *vcf*. We ran additional basic filtering as recommended by the tool developers to filter to high-confidence variants. Specifically, we first generated a pileup file with *samtools mpileup* followed by *snpfilter.pl* with default settings. Followed by *prepare_for_readcount.pl* and *bam-readcount* to calculate the number of reads mapping at each position. Finally, we ran *fpfilter.pl and highconfidence.pl* to run the false positive filter and filter based on likelihood of being a somatic variant and mapping quality. To further remove likely false positives, we also filtered out variants with <8 reads mapping to the locus in the normal BAM file and <5 reads mapping to the Alt allele in the tumour BAM file.

**Mutect2**

Mutect2[5] (v4.6.1.0) was run on both the linear and pangenome aligned BAM files using GATK Muect2 and the following parameters: -normal-sample, -reference.fa, -intervals.list, and two -input BAM files, *tumour.bam* and *normal.bam* samples. To annotate variant allele frequencies, bcftools +fill-tags was applied to populate the INFO-field AF tag. To remove likely false positives, we ran FilterMutectCalls by first generating pileup for both the tumour and normal samples using GetPileupSummaries, then calculated cross-sample contamination using CalculateContamination. Only variants flagged as “PASS” were considered. We filtered out INDELs and kept SNVs only. To further remove likely false positives, we also filtered out variants with <8 reads mapping to the locus in the normal BAM file and <5 reads mapping to the Alt allele in the tumour BAM file.

**Consensus SNV detection**

Previous benchmarking of somatic SNV detection using the linear reference genome demonstrated that ensemble-based methods, where SNVs were only considered if called by multiple tools, outperformed somatic SNV detection by a single tool. Thus, for each individual, we developed a consensus SNV call set considering only SNVs that were detected by two or more tools.

**Somatic SNV Detection Performance**

For each tool, performance was evaluated against somatic SNV calls from the TCGA MC3 project[6]. Comparisons were done in GRCh38 coordinates and liftover across assemblies were done using Picard LiftoverVcf (v3.3.0). SNVs were determined as a true positive if the chromosome, position and alt allele matched that reported by the MC3. SNVs called here but not by the MC3 were considered false positives. SNVs called by MC3 but not here were considered false negatives. Using these definitions, we calculated the precision, recall and F1-score for each tool with each reference genome against the TCGA MC3 gold standard.

**Evaluating Germline Contamination**

We considered all SNVs detected by Strelka2 when aligned to the pangenome as this was the highest performing combination. Putative germline variants were determined from gnomAD[7] (v2).

We considered all variants reported in gnomAD as well as only common variants, defined as minor allele frequency (MAF) > 0.01. We calculated the number of somatic SNVs that overlapped germline variants reported in gnomAD using bedtools closest (v2.30.0) defining an overlap if distance = 0. We evaluated the number of germline-somatic overlaps when aligning to the pangenome vs linear reference and across ancestry populations. Differences in germline-somatic overlap were quantified using Mann-Whitney Test.

#### Evaluating Reference Bias

To evaluate reference bias, at each locus, we calculated the number of reads that map to the reference allele vs the alternate allele. We considered the alignment as showing high reference bias if the number of reads mapping to the reference allele far exceeded the number of reads mapping to the alternate allele. We considered only the somatic SNVs detected by Strelka2 when aligned to the pangenome as this was the highest performing combination. The number of reads that mapped to the alternate and reference alleles were extracted from the Strelka2 VCF. However, because an imbalance in read counts mapping to the reference vs alternate allele would also be expected in the case of subclonal variants, we also considered heterozygous germline variants in close proximity to the somatic SNVs. We considered heterozygous germline variants within +/- 75bp (the average read length) and similarly calculated the number of reads mapping to the reference vs alternate allele using GATK HaplotypeCaller. **Supplementary Figure 7a** shows the number of proximal heterozygous germline variants for each donor used in this analysis.

#### Evaluating consequences of MC3-excusive and pangenome-exclusive SNVs

We defined pangenome-exclusive SNVs as somatic SNVs identified leveraging Strelka2 with alignment to the pangenome but not found by TCGA MC3. We defined MC3-exclusive SNVs as somatic SNVs identified by TCGA MC3 but not Strelka2 with pangenome alignment. Next, we leveraged definitions of cancer driver genes by Bailey et al.[8] and categorized genes as bladder cancer specific, pancancer or both bladder and pancancer. We quantified the number of MC3-exclusive and pangenome-exclusive SNVs within each category of driver gene, further stratifying variants based on predicted impact by SnpEff[9].

#### Replication in lung adenocarcinoma

We selected 29 lung adenocarcinoma patients from The Cancer Genome Atlas (TCGA) based on inferred genetic ancestry: 10 of African ancestry, 9 of East Asian ancestry (only 9 were available) and 10 of European ancestry according to Yuan *et al.*[1]. Individuals were selected based on maximum proportion of each ancestry population. Whole exome sequencing BAM files for both tumour and matched normal aligned to GRCh38 were downloaded from GDC. Reads were back-extracted from BAM files using samtools bamtofastq (v 1.16.1). Tumour and blood whole exome sequencing was aligned to the human pangenome reference (v1.1) as described above. Strelka2[3] (v2.9.10) was run on both the linear and pangenome aligned BAM files using the somatic workflow configuration script configureStrelkaSomaticWorkflow.py as described above.

Performance was evaluated against somatic SNV calls from the TCGA MC3 project[6] as described above. There were three tumours with F1-score < 20% when aligned to both the linear and pangenome references due to few (< 10) SNVs detected by MC3 resulting in low precision with both alignments.

**Data Visualization**

The graphs were generated using R version 4.4.2 with ggplot2 version 4.4.0.
